# Dual targeting of KDM1A and antioxidants is an effective anticancer strategy

**DOI:** 10.1101/2024.06.12.597953

**Authors:** Shaila Mudambi, Megan E Fitzgerald, Deschana L Washington, Paula J Pera, Wendy J Huss, Gyorgy Paragh

**Affiliations:** Department of Cell Stress Biology, Roswell Park Cancer Institute, Elm and Carlton Streets, Buffalo, NY, United States 14263; Department of Dermatology, Roswell Park Cancer Institute, Elm and Carlton Streets, Buffalo, NY, United States 14263

**Keywords:** cancer, reactive oxygen species, KDM1A, HIF-1A, antioxidants

## Abstract

Lysine Specific Demethylase 1 (KDM1A / LSD1) regulates mitochondrial respiration and stabilizes HIF-1A (hypoxia-inducible factor 1A). HIF-1A modulates reactive oxygen species (ROS) levels by increasing cellular glucose uptake, glycolysis, and endogenous antioxidants. The role of KDM1A in cellular ROS response has not previously been described. We determined the role of KDM1A in regulating the ROS response and the utility of KDM1A inhibitors in combination with ROS-inducing cancer therapies. Our results show that KDM1A inhibition sensitized cells to oxidative stress and increased total cellular ROS, which was mitigated by treatment with the antioxidant N-acetyl cysteine. KDM1A inhibition decreased basal mitochondrial respiration and impaired induction of HIF-1A after ROS exposure. Overexpression of HIF-1A salvaged cells from KDM1A inhibition enhanced sensitivity to ROS. Thus we found that increased sensitivity of ROS after KDM1A inhibition was mediated by HIF-1A and depletion of endogenous glutathione. We also show that KDM1A-specific inhibitor bizine synergized with antioxidant-depleting therapies, buthionine sulfoximine, and auranofin in rhabdomyosarcoma cell lines (Rh28 and Rh30). In this study, we describe a novel role for KDM1A in regulating HIF- 1A functions under oxidative stress and found that dual targeting of KDM1A and antioxidant systems may serve as an effective combination anticancer strategy.

## Introduction

Elevated levels of reactive oxygen species (ROS), redox imbalance, and deregulation of redox signaling pathways are essential hallmarks of cancer(1). Cancer cells persistently show increased levels of ROS. ROS-induced DNA damage can cause the upregulation of oncogenes or inactivation of tumor suppressors contributing to tumor progression(2). Increased hydrogen peroxide levels stimulate the constitutive activity of multiple kinases, including c-jun(3–5), c-fos(5), MAPKs(6,7), and enhance growth factor receptor signaling through the EGF and PDGF pathways(8,9). Thus, hydrogen peroxide is essential for cell proliferation in multiple cancers.

However, cancer cells not only elevate hydrogen peroxide levels to sustain signaling but also upregulate antioxidant activity to maintain the balance between ROS generation and elimination and evade the detrimental effects of excess ROS(10). To maintain redox balance, numerous cancers upregulate the antioxidant machinery and increase the levels of endogenous antioxidants like glutathione and thioredoxin (11).

Lysine specific histone demethylase (LSD1/KDM1A) is a transcriptional coregulator that controls transcription through a FAD-dependent enzymatic reaction which results in the removal of mono- or di-methyl groups from both histone and non-histone proteins.(12,13) KDM1A has been shown to regulate cellular redox state by modifying mitochondrial activity(14,15) as well as by stabilizing proteins involved in metabolic adaptation like HIF-1A protein(16), and p53(17–19), E2F1(20–22) and myc(23,24). High expression of KDM1A has been linked to poor prognosis in cancer patients.(25–29) Furthermore, KDM1A inhibitors are used to target epigenetic gene regulation in cancer and have been tested in clinical trials but have had limited success(30,31). We have also shown previously that treatment with KDM1A inhibitor greatly improved the efficacy of photodynamic therapy(32).

Although there is growing evidence that KDM1A plays a role in redox regulation, the link between KDM1A and cellular ROS levels in cancer has not been studied. The current studies show that KDM1A plays a role in regulating cellular ROS via HIF-1A and show that a novel approach of targeting KDM1A to sensitize cells may improve anticancer efficacy of exogenous ROS inducing cancer therapy agents.

## Material and methods

### Cell Culture

All cells were maintained in humidified 5% CO_2_ incubator at 37 °C. Cell lines (Table 1) were cultured in either complete media - DMEM or MEM (Corning, NY, USA) supplemented with 1mM Corning glutagro supplement (Corning, NY, USA), 1% antibiotic solution (100 mg/L streptomycin, 100 U/ml penicillin) and 10% fetal bovine serum (Corning, NY USA). Cell lines used are described in Table 1.

**Table 1:** Cell Lines.

| Cell Line | Type | Source |
| --- | --- | --- |
| **HaCaT** | human, adult, low calcium, high temperature<br>keratinocytes cells | Addex Biosciences |
| **FaDu** | Human head and neck squamous cell carcinoma | ATCC |
| **Rh28** | Human Rhabdomyosarcoma cells | ATCC |
| **Rh30** | Human Rhabdomyosarcoma cells | ATCC |

### Transfection and transduction

Cells were trypsinized with 0.25% trypsin (Corning, NY, USA) and plated in Costar 6 well (2*10^5^ cells/well) or Costar 100mm plates (1*10^6^ cells per plate) (Corning, NY, USA).

Transfection with either plasmids (2-10 μg) or siRNA (100 nM) was carried out using JetPrime transfection reagent (PolyPlus transfection, USA) according to manufacturer instructions as cells attached to the plate overnight. Next day transfection media was removed. 48 hours after transfection cells were plated and all treatments for transiently transfected cells were performed within 72-96 hours.

Viruses were prepared using the 293T cell line (Clontech, USA) transfected with the appropriate expression vector along with packaging plasmids(ΔR) and G-protein vesicular stomatitis virus (VSVG) for lentivirus and gag-pol for retrovirus using JetPrime transfection reagent (PolyPlus transfection, USA) according to manufacturer instructions. Used plasmids are listed in Table 2. Viral supernatants were collected 48h and 72h later, filtered through disposable 0.45 μM cellulose acetate filters and frozen in individual aliquots at -80°C. For infection cells were plated in 60- or 100-mm tissue culture dishes and allowed to achieve 60% confluence before adding viral supernatant in the presence of 8 μg/ml polybrene for 24 hours (Sigma, St. Louis, MO). Cells were selected by treatment with puromycin (1ug/ml for FaDu cells and 2ug/ml for HaCaT) for 5 days.

**Table 2:** Plasmids and siRNA.

| Reagent | Source |
| --- | --- |
| PLVP- Flag plasmid | Nikiforov Lab(33) |
| Flag HIF-1A | Semenza Lab(34) |
| siGENOME Non-Targeting siRNA Pool #2 | Cat # D-001206-14-05 Dharmacon |
| SMARTpool: siGENOME KDM1A siRNA | Cat # M-009223-01 Dharmacon |

### Cell treatment

#### N-acetylcysteine (NAC)

Cells were trypsinized and plated in 96 wells (2500 cells per well) or 6 well plates (1*10^5^ cells per well). All drug treatments were performed 24 hours prior to hydrogen peroxide exposure. Stock concentration of hydrogen peroxide (Sigma, St. Louis, MO) was made in 1X PBS and added to plates to make the final working concentration. The pH of NAC when used was adjusted to 7.4 with sodium bicarbonate before adding to cells. Cells were treated with 10mM NAC (Sigma, St. Louis, MO) and bizine for 18 hours prior to hydrogen peroxide exposure.

#### Drug treatments

Cells were plated at 2500-3000 cells per well in 96 well plates, bizine and tested drug was added simultaneously to the wells, and cell viability or survival was assessed, according to manufacturer’s protocol, 24-48 hours after the addition of bizine and tested drug (Table 4).

### Cell viability assays

Cell viability was measured using cell titer blue assay or methylene blue survival assay 24 hours post hydrogen peroxide exposure. 6X Cell Titer Blue solution (Promega Corporation, Madison, WI) was added into wells with cells and wells with plain media as a no cell control, to a final 1X concentration and incubated for 3 hours at 37℃ and 5% CO_2._ Fluorescence was measured using a 560 nm excitation/590nm emission filter set Cell viability with cell titer blue is based on metabolic conversion of resorufin and therefore methylene blue survival assay was used to determine cell death in Figure 5 since HIF-1A modifies cellular metabolism. Cells were incubated with 0.5% methylene blue in 50% Methanol (Sigma, St. Lois, MO) for 30 min and excess methylene blue was washed with double distilled water and dried. Methylene blue was dissolved in 500µl of 1% sodium dodecyl sulfate (SDS) solution (Corning, NY) and 100µl was transferred into a 96 wells plate and measured at 540 nm.

### ROS measurements using flow cytometry

#### Total cellular ROS using Cell ROX dye

Cells were plated in 6 well plates at 1.5*10^5^ per well. Next day cells were treated with bizine or phenelzine. 24 hours after drug treatment, cells were exposed to hydrogen peroxide for 5 min. Hydrogen peroxide was added in 1X PBS with calcium (200 mg/L) and magnesium (98 mg/L) supplemented with 25mM glucose. Wells were washed once with 1X PBS and 1 mL of complete media was added back to well. Cell ROX dye (Thermofisher Scientific, USA) was added to a final concentration of 10μM. The dye was incubated with cells for 30 min at 37℃ and 5% CO_2_. Media containing cell ROX was removed and wells were washed twice with 1X PBS, trypsinized and harvested into flow tubes. Cells were pelleted at 200 g for 10 min and resuspended in 300ul of 1X PBS for analysis by flow cytometry.

#### Lipid peroxidation using BODIPY

Cells were plated in 6 well plates at 1.5*10^5^ per well. 24 hours post plating cells were treated with bizine. 24 hours after treatment hydrogen peroxide added in complete media for 90min. Wells were washed once with 1 ml of 1x PBS with glucose and magnesium and calcium and added back to wells. BODIPY® 581/591 C11 reagent was added to a final concentration of 10μM. The dye was incubated with cells for 30 min at 37℃ and 5% CO_2_. Cells treated with cumene hydroperoxide for 1 hour was used as positive control. Media containing BODIPY® 581/591 C11 reagent (Thermofisher Scientific, USA) was removed and wells were washed twice with 1X PBS, trypsinized and harvested into flow tubes. Cells were pelleted at 200 g for 10 min and resuspended in 300µl of 1X PBS for analysis by flow cytometry.

### Quantitative PCR

Total RNA was isolated using RNeasy Mini Kit (Qiagen Inc, MD, USA) according to the manufacturer’s protocol. cDNA was synthesized from 500 ng total RNA using High-Capacity cDNA Reverse Transcription Kit (Thermofisher Scientific, USA) in a 20μl reaction according to the manufacturer’s protocol. Quantitative reverse-transcription PCR was performed using QS6 PCR System (Applied Biosystems, Carlsbad, CA, USA) using SYBR Green Master Mix (Thermofisher Scientific, USA) or TaqMan Universal Master Mix II (Applied Biosystems, Carlsbad, CA, USA). All reactions were performed in triplicate, and the experiments were repeated at least twice. The results are presented as the mean of at least 2 experiments. Sybr green primers used: GLRX-1 (F- GATTGGAGCTCTGCAGTAACCA, R- CAATGCCATCCAGCTCTTGA), HMOX-1 (F- TTCTCCGATGGGTCCTTACACT, R- GGCATAAAGCCCTACAGCAACT), KLF-9 (F- CTCCGAAAAGAGGCACAAGT, R- CGGGAGAACTTTTTAAGGCAGT), SDHA (F- TGGGAACAAGAGGGCATCTG, R- CCACCACTGCATCAAATTCATG) and PPIA (F- TTATTTGGGTTGCTCCCTTC, R- AAGTGTGCCAAATCTGCAAG). Taqman probe: KDM1A (Cat # #4331182 ThermoFischer).

### Immunoblotting

Protein extracts were prepared by lysing cells using 1x lysis buffer (1% SDS, 0.01% Tris- Hcl). Protein concentration was estimated using Pierce™ BCA Protein Assay Kit (Thermofisher Scientific, USA). Equal amounts of protein were run on 4-20% gradient precast gels and transferred to PVDF membrane. Membranes were blocked with 5% milk and incubated with primary antibodies overnight and incubated with anti-rabbit secondary antibody from Cell Signaling Technologies for one hour after which blots were developed using Bio-Rad ChemiDoc™ Imaging Systems. All antibodies (Table 3) were purchased from Cell Signaling Technologies.

**Table 3:**
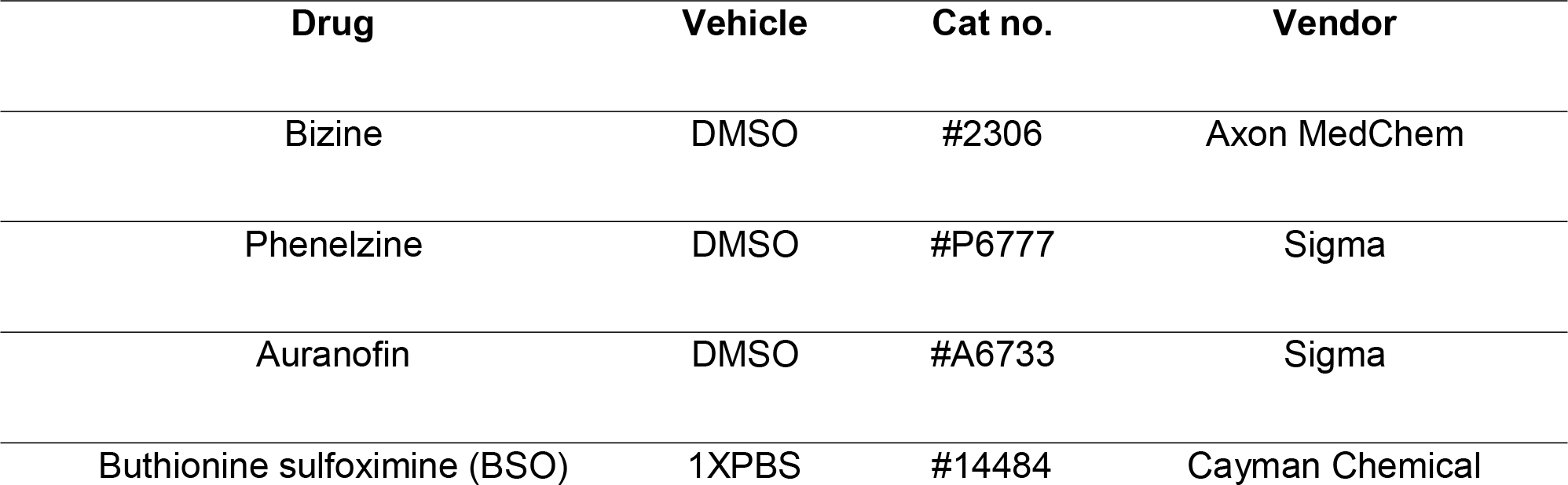
List of drugs used.

| **Drug** | **Vehicle** | **Cat no.** | **Vendor** |
| --- | --- | --- | --- |
| Bizine | DMSO | #2306 | Axon MedChem |
| Phenelzine | DMSO | #P6777 | Sigma |
| Auranofin | DMSO | #A6733 | Sigma |
| Buthionine sulfoximine (BSO) | 1XPBS | #14484 | Cayman Chemical |

**Table 4:** List of Antibodies used.

| **Target Protein** | **Cat no.** | **Type** | **Dilution** |
| --- | --- | --- | --- |
| **KDM1A** | 2184 | Rabbit<br>monoclonal | 1:1000 |
| **HIF-1A** | 14179 | Rabbit<br>monoclonal | 1:1000 |
| **Lamin A/C** | 2032 | Rabbit<br>monoclonal | 1:3000 |
| **β- Tubulin** | 2146 | Rabbit polyclonal | 1:2000 |

### Seahorse assay

Investigation of oxygen consumption rate/ extracellular acidification rate (OCR/ECAR) ratios and mitochondrial function were performed using XF96 analyzers with XF96 V3 cell culture microplates (Agilent, Santa Clara, CA, USA). FaDu and HaCaT cells were plated at 8000 cells per well and treated with bizine 24 hours after plating. Next day OCR was assessed using Seahorse assay buffer (containing 10 mM glucose, 10 mM pyruvate, pH 7.4) or basic glucose free DMEM medium (pH 7.4) respectively. The following compounds and concentrations were added dependent on type of experiment: glucose (25 mM); oligomycin A (1 μM); FCCP (optimized concentrations; 1uM for both HaCaT and FaDu); antimycin A (2.5 μM); rotenone (2.5 μM); 2-deoxyglucose (10 mM or 50 mM). Readings were normalized to OD of cells growing in parallel plates, treated the same and stained with methylene blue.

### Glutathione level measurement

Cells were plated at 2500 cells per well in a 96 well and treated with bizine the next day. 2 hours after bizine treatment cells were exposed to 250μM or 500μM hydrogen peroxide for 2 min after which glutathione (GSH) levels in cells were assayed according to the manufacturer’s instructions for GSH-Glo™ Glutathione Assay (Promega, Madison, WI, USA).

### Statistics and Data Analysis

Data analysis was performed using GraphPad Prism Version 9.5.1 . Significance was calculated where applicable using student t-test.

## Results

### KDM1A inhibition sensitizes cells to oxidative stress

To determine the effects of KDM1A on regulation of ROS response, cell viability was assessed in FaDu and HaCaT cells by chemically inhibiting KDM1A with increasing doses of KDM1A specific inhibitors, bizine (Figure 1A, C) and phenelzine (Figure 1B, D) for 24 hours and then exposing cells to hydrogen peroxide. Treatment with inhibitors significantly decreased cell viability in both cell lines to hydrogen peroxide induced oxidative stress even at inhibitor doses that were non-toxic to cells. To rule out non-KDM1A specific effects of the inhibitor, expression of KDM1A was knocked down using siRNA in both HaCaT and FaDu cells (Figure 1E-H) and then cells were exposed to hydrogen peroxide. KDM1A inhibition by siRNA sensitized both cell lines to oxidative stress in a dose dependent manner (Figure 1 F, H). Further, to investigate whether treatment with bizine and hydrogen peroxide had any effect on KDM1A protein expression, FaDu cells were treated with bizine and then exposed to hydrogen peroxide.

**Figure 1:**
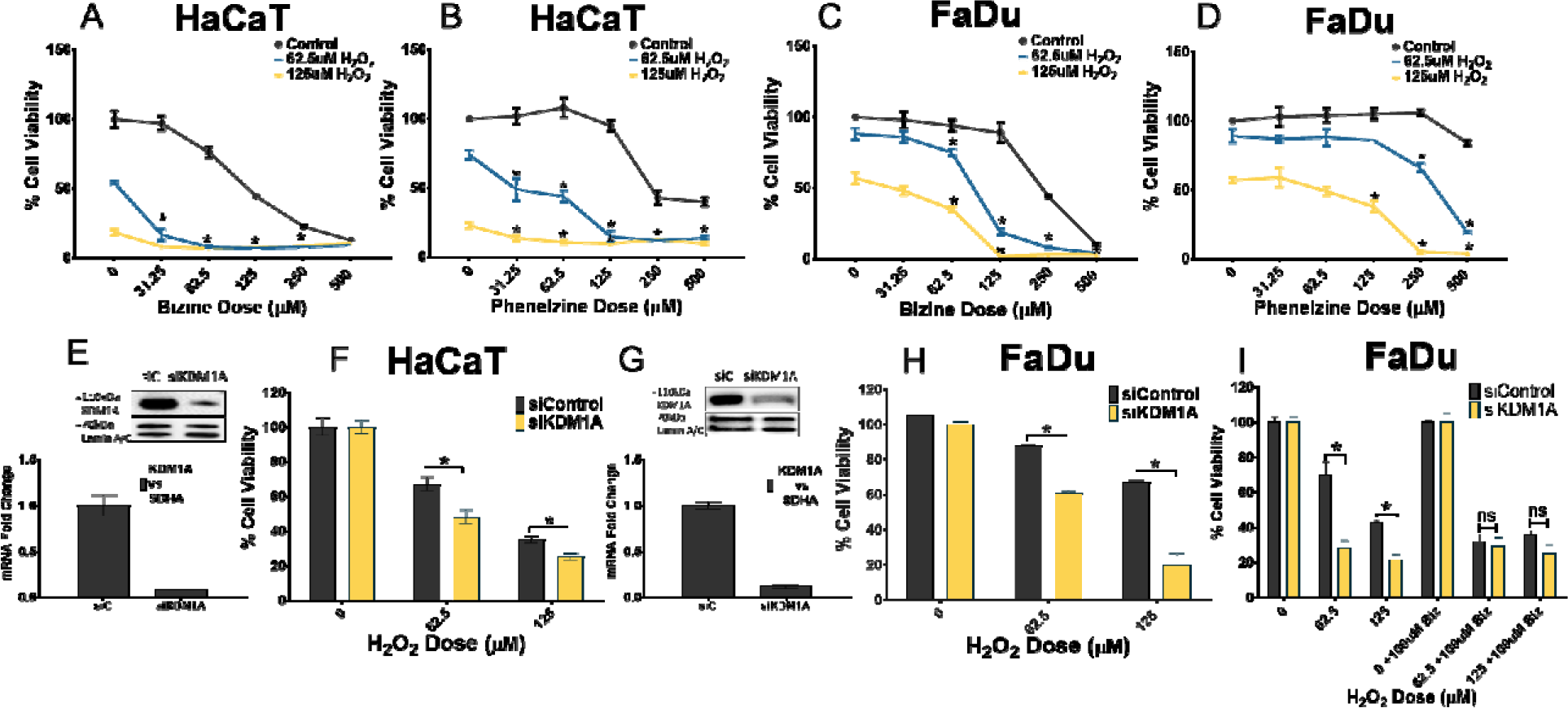
KDM1A inhibition sensitizes cells to oxidative stress: Cell viability in HaCaT and FaDu cells treated with (A, C) 0-500µM of bizine and (B, D) 0-500µM phenelzine for 24 hours after exposure to hydrogen peroxide. (E, G) The mRNA and the protein levels of KDM1A 48hr after KDM1A downregulation using siRNA in HaCaT and FaDu cells (siC- Control siRNA, siKDM1A – KDM1A siRNA). Values are expressed as fold change over control. (F, H) Effect of KDM1A downregulation on cell viability after hydrogen peroxide treatment in HaCaT and FaDu cells. Cell viability was assessed using cell titer blue assay and is expressed as a percentage of 0 µM H_2_O_2_ or siControl as appropriate. Bars represent mean ± SD (n=3). *: p<0.05.

Treatment with bizine or hydrogen peroxide did not have a significant impact on KDM1A protein expression (Figure S1)

### Effects of KDM1A inhibition on ROS response is mediated by increase in cellular ROS levels

Building on our findings that KDM1A plays a role in regulating cellular redox balance, we further investigated the impact of KDM1A inhibition on the levels of reactive oxygen species (ROS) in FaDu cells. Specifically, changes in total cellular ROS levels and lipid peroxidation were assessed using flow cytometry under baseline conditions and after treatment with hydrogen peroxide, with or without KDM1A inhibition (Figure 2). There was no observable difference in cellular ROS or lipid peroxidation in FaDu cells after bizine treatment until after the addition of hydrogen peroxide. Bizine treatment combined with hydrogen peroxide significantly increased the levels of total cellular ROS (measured by cell ROX) (Figure 2A, B). Additionally, FaDu cells treated with bizine, or with KDM1A knockdown, demonstrated a dose-dependent increase of peroxidation of lipids after exposure to hydrogen peroxide (Figure 2C, D). We also assessed the expression of genes commonly upregulated in response to oxidative stress using qPCR post bizine and hydrogen peroxide treatment. Out of the six genes tested, only KLF-9 and glutaredoxin 1 (GLRX-1) expression was upregulated after hydrogen peroxide exposure in bizine treated (Figure 2E, F) or in KDM1A downregulated (Figure 2G, H) cells. In bizine treated cells GLRX-1 levels were upregulated at baseline but not in the highest dose of treatment. This was observed in multiple experiments. It has been reported that an increase in ROS beyond a certain threshold will induce the degradation of GLRX-1 through caspase 3 and 9(35). It is possible that GLRX-1 is regulated by a similar mechanism here and that the ROS levels at the highest bizine and hydrogen peroxide dose reaches the threshold level. Interestingly, several other antioxidant genes tested including TXNRD1, SOD1, CAT and NQO1 remained unchanged (Figure S2). To confirm whether the increase in ROS was directly mediating cell death in our system, a rescue experiment using antioxidant N-acetylcysteine (NAC) was performed. FaDu cells were treated with bizine and NAC 18 hours prior to hydrogen peroxide exposure.

**Figure 2:**
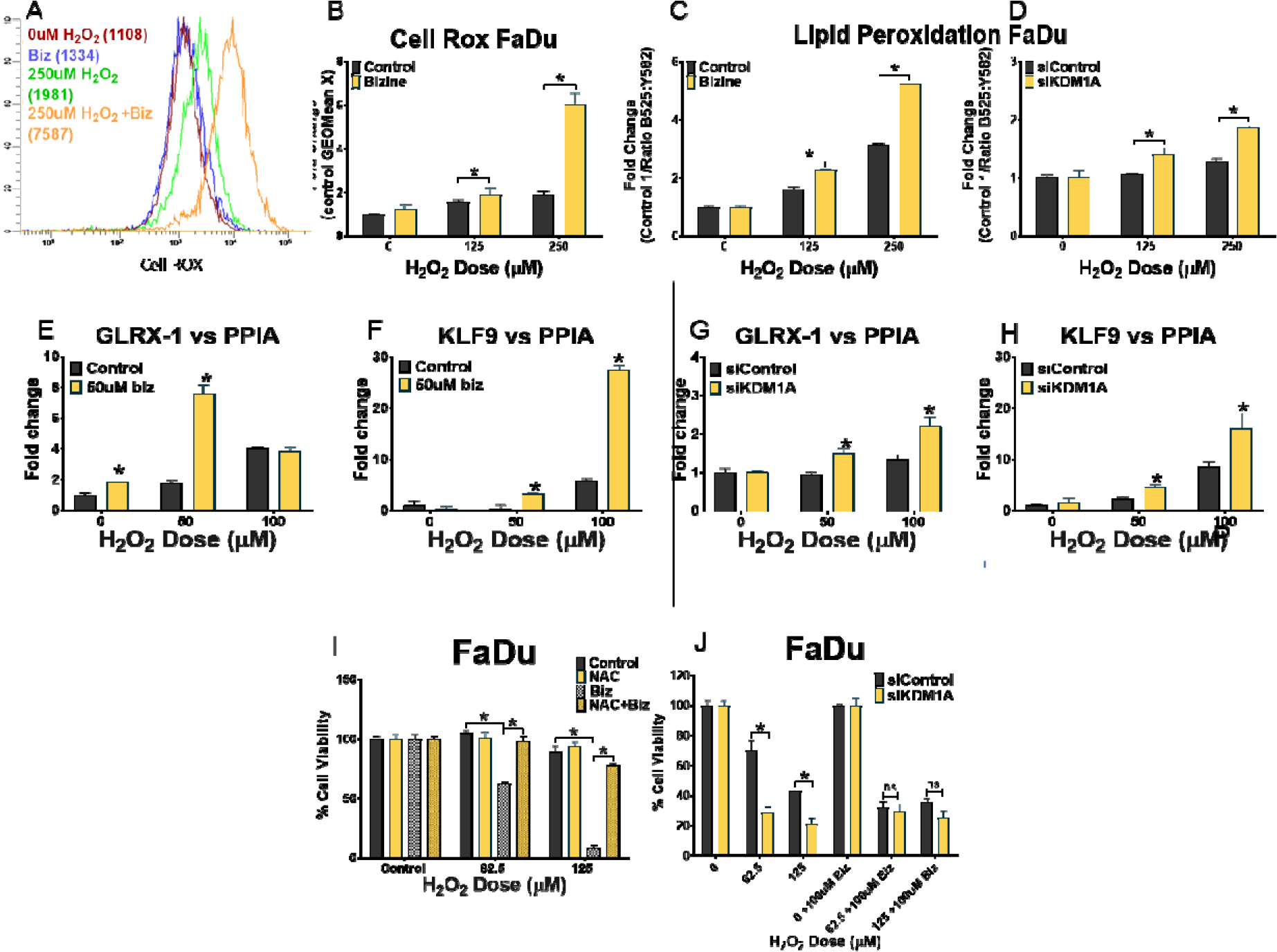
Effects of KDM1A inhibition on ROS response is mediated by increase in cellular ROS levels: (A) Total cellular ROS in FaDu cells treated with 150µM of bizine for 24 hours and 5 min of hydrogen peroxide. (B) Fold change of cellular ROS as measured by Cell ROX dye in FaDu cells treated with 150µM of bizine for 24 hours and 5 min of hydrogen peroxide. (C, D) Lipid peroxidation by staining with BODIPY® 581/591 C11 in FaDu cells treated with 150µM of bizine for 24 hours (C) or in cells with KDM1A downregulated (D) after 90min of hydrogen peroxide treatment. (E-H) Gene expression of ROS regulated genes (GLRX-1 and KLF-9) after 24 hours of 50µM bizine treatment (E, F) or after KDM1A inhibition (G, H) and hydrogen peroxide exposure. RNA was harvested 4 hours after the addition of hydrogen peroxide in FaDu cells. Values are expressed as a fold change over control (I) Cell viability in FaDu cells after 18 hours of 150uM of bizine and 10mM NAC treatment and hydrogen peroxide exposure. (J) Cell viability in bizine treated FaDu cells after KDM1A downregulation and hydrogen peroxide treatment. Cell viability was assessed using cell titer blue assay and is expressed as a percentage of control. Bars represent mean ± SD (n=3). *: p<0.05.

Treatment with the NAC rescued cells from enhanced sensitivity to ROS resulting from bizine treatment (Figure 2I), indicating that NAC was able to mitigate the effects of increased ROS levels caused by KDM1A inhibition. The same effect was seen in HaCaT cells (Figure S3). The combination of KDM1A siRNA knockdown and bizine treatment had no additional effect on cell viability in FaDu cells, suggesting that KDM1A is needed for bizine to exert its effects (Figure 2J).

### KDM1A inhibition modifies mitochondrial energetics; decreases basal respiration, mitochondrial ATP generation and cellular glutathione levels

To determine if increased cellular ROS was responsible for cell death when combined with KDM1A inhibition the seahorse assay was used to assess mitochondrial activity with KDM1A inhibition. Bizine treatment decreased basal respiration rate and ATP production by the mitochondria in FaDu cells (Figure 3 A, B) and in HaCaT cells (Figure S4). KDM1A inhibition did not enhance the activity of the most prominent source of ROS, the mitochondria(36). Bizine treatment reduced the ratio of GSH to GSSG in a dose dependent manner compared to control (Figure 3C). KDM1A was downregulated using siRNA and showed a decrease in the ratio of GSH and GSSG similar to bizine treatment (Figure 3C). Importantly the decrease in ratio is observed at baseline even prior to the addition of hydrogen peroxide suggesting that the absence of KDM1A results in impairment in glutathione production.

**Figure 3:**
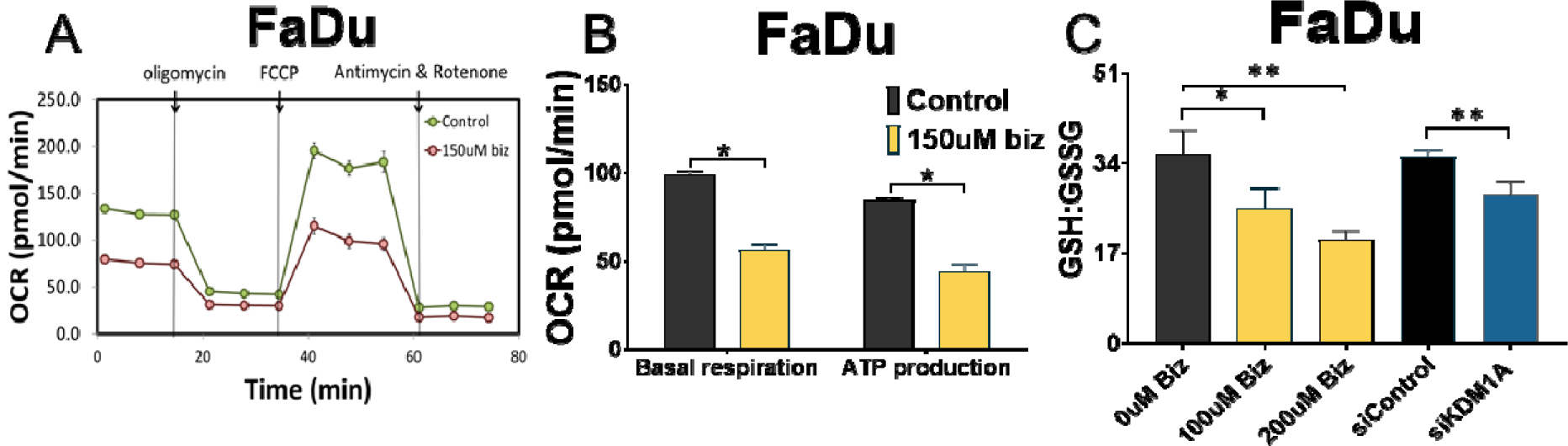
Effects of KDM1A inhibition on mitochondrial energetics and cellular glutathione levels: (A) MitoStress test using seahorse assay in FaDu cells after bizine treatment. (B) Graphs quantifying basal respiration and ATP generation in bizine treated FaDu cells. (C) Ratio of GSH and GSSG after 2 min of 500µM hydrogen peroxide treatment in FaDu cells after KDM1A downregulation with siRNA. Bars represent mean ± SD (n=3). *: p<0.1, **p<0.05.

### KDM1A effects on ROS regulation are mediated by impairment of HIF-1A induction

HIF-1A is a regulator of cellular metabolic pathways and plays a role in maintaining cellular redox status and glutathione levels by regulating antioxidant enzymes including GLRX-1 and mitochondrial activity(37). KDM1A has been shown to regulate the stability and activity of HIF-1A post translationally by multiple mechanisms(16,38–40). To assess whether HIF-1A mediated changes plays a role in KDM1A mediated ROS response, we determined the effect of KDM1A inhibition on HIF-1A protein expression in HaCaT and FaDu cells. Bizine treatment reduced, but did not completely inhibit, the induction of HIF-1A protein after hydrogen peroxide treatment (Figure 4A-D). To assess the functional role of HIF-1A in KDM1A mediated ROS response in both cell lines, HIF-1A was downregulated in FaDu cells using two different lentiviral shRNA vectors. Downregulation of HIF-1A was validated by exposing cells to hydrogen peroxide (Figure 4E). FaDu cells were treated with bizine and exposed to hydrogen peroxide.

**Figure 4:**
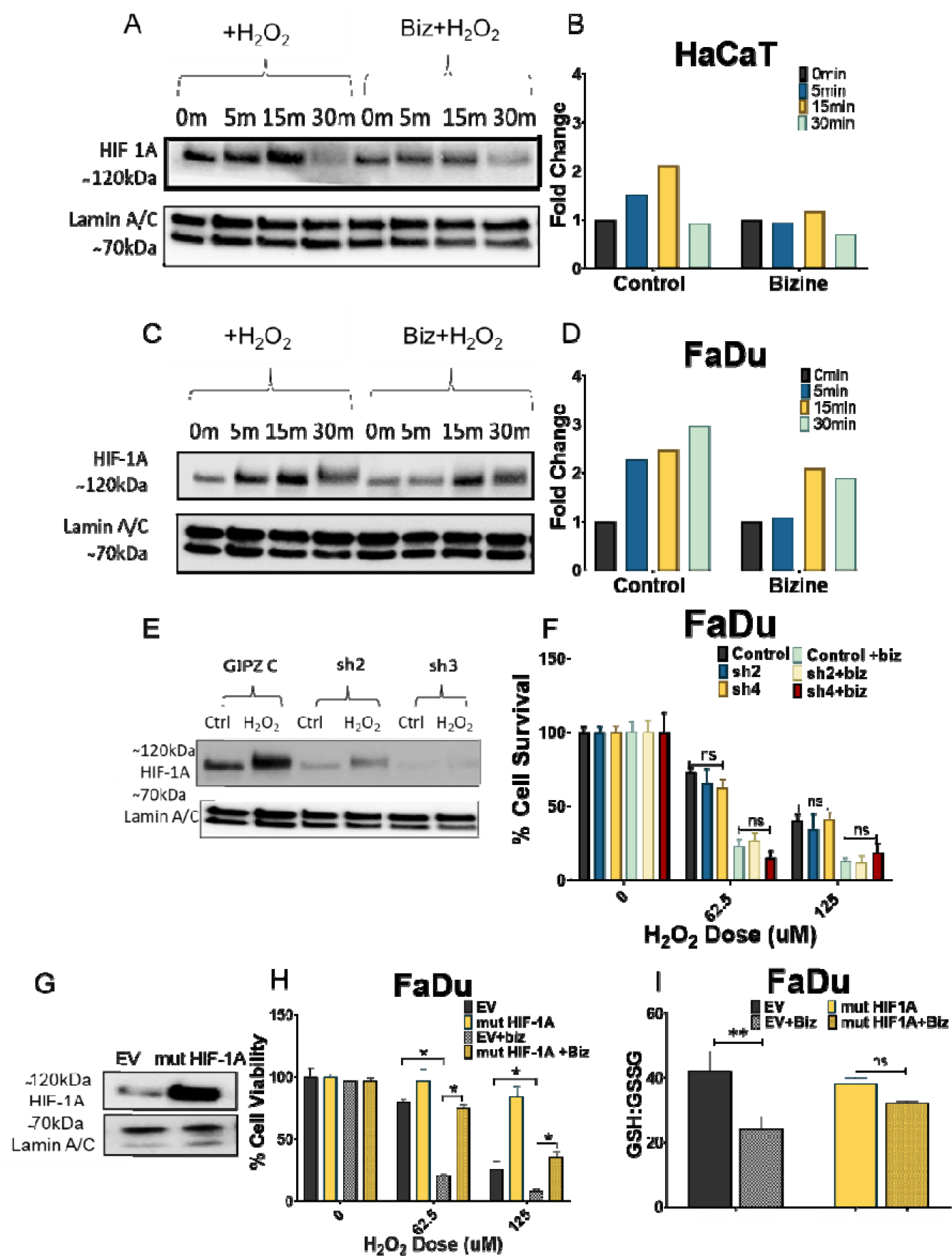
**KDM1A inhibition effects on ROS regulation are mediated by impairment of HIF- 1A induction:**(A-D) HIF-1A protein levels at 0-30 minutes after bizine and 500µM hydrogen peroxide treatment in HaCaT and FaDu cells (Western blot). (E) HIF-1A lentiviral shRNAs; sh1 and sh2 downregulates HIF-1A protein in control cells and after 15 min of hydrogen peroxide treatment (Western blot). (F) Cell survival measured using methylene blue assay 24 hours of bizine and 5 min of hydrogen peroxide treatment in FaDu cells with HIF-1A downregulated using 2 different shRNA; sh1 and sh2. (G) Constitutively active HIF-1A mutant is overexpressed in FaDu cells (Western blot; EV: Empty vector; Mut HIF1A: constitutively active HIF1A variant overexpressed). (H) Cell viability measured by cell titer blue assay after 24 hours of bizine and 15 min hydrogen peroxide treatment in cells expressing constitutively active HIF-1A mutant (I) Ratio of reduced (GSH) and oxidized (GSSG) glutathione levels in FaDu cells treated for 24 hours with bizine after HIF-1A constitutively active mutant overexpression (EV: Empty vector; Mut HIF1A: constitutively active HIF1A variant overexpressed). Bars represent mean ± SD (n=3). **: p<0.005, *<p<0.05.

HIF-1A inhibition did not decrease cell viability at baseline after bizine treatment (Figure 4F) indicating that HIF-1A may be partially mediating effects of KDM1A inhibition. A constitutively active mutant HIF-1A was overexpressed in FaDu cells to determine response to hydrogen peroxide after bizine treatment. Overexpression of HIF-1A salvaged cells from bizine enhanced hydrogen peroxide sensitivity (Figure 4H). HIF-1A has also been shown to increase the levels of glutathione in cells(37), therefore we determined levels of reduced GSH and oxidized glutathione GSSG in FaDu cells with overexpressed HIF-1A treated with bizine and hydrogen peroxide. Levels of GSH and GSSG were maintained in bizine treated cells with HIF-1A constitutively expressed (Figure 4I) indicating that the impairment of HIF-1A induction is responsible for mediating effects of KDM1A inhibition on oxidative stress.

### Inhibitors of antioxidant generation, auranofin and buthionine sulfoximine synergize with bizine to enhance cell death in rhabdomyosarcoma cells

Since KDM1A inhibition impaired cellular antioxidant capacity, KDM1A inhibitors were tested in combination with chemotherapies that induce ROS; cisplatin and elesclomol or with other antioxidant inhibitors; auranofin and Buthionine sulfoximine (BSO). Cisplatin has been shown to induce mitochondrial ROS in squamous carcinoma cell lines(41). Therefore, cell viability was tested when FaDu, HaCaT and A-431 squamous cell carcinoma cells were treated with a combination of cisplatin and bizine. Bizine was incapable of enhancing the effect of cisplatin in all tested cells lines (Figure 5A, E, I). Elesclomol has been shown to interact with the electron transport chain to generate ROS in melanoma cells(42). Cell viability was tested when FaDu cells and B16-F10 and MNT-1 melanoma cells were treated with a combination of elesclomol and bizine. Bizine was incapable of enhancing the effect of elesclomol in the tested cells lines (Figure 5B, F, J). Auranofin and BSO are drugs that target the two major cellular antioxidant pathways; thioredoxin and glutathione (43). Auranofin is a thioredoxin reductase inhibitor that has been demonstrated to elevate reactive oxygen species (ROS) levels in various cell lines (44). On the other hand, BSO inhibits the enzyme responsible for the rate-limiting step in glutathione synthesis, leading to increased ROS within cells as a result of depleted cellular glutathione levels (45). Additionally, BSO has been used in combination with auranofin for rhabdomyosarcoma (43). Therefore, we tested the cell viability of not only FaDu but also Rh28 and Rh30 rhabdomyosarcoma cells when treated with bizine and auranofin (Figure 5 C, G and K) or bizine and BSO (Figure D, H and L). Bizine increased auranofin induced cell death in FaDu, Rh28, and Rh30 (Figure 5 C, G and K). Bizine synergized with BSO showing a coeffecient of drug interaction (CDI)<1 in enhancing cell death in the rhabdomyosarcoma cell lines (Figure 5 H and L).

**Figure 5:**
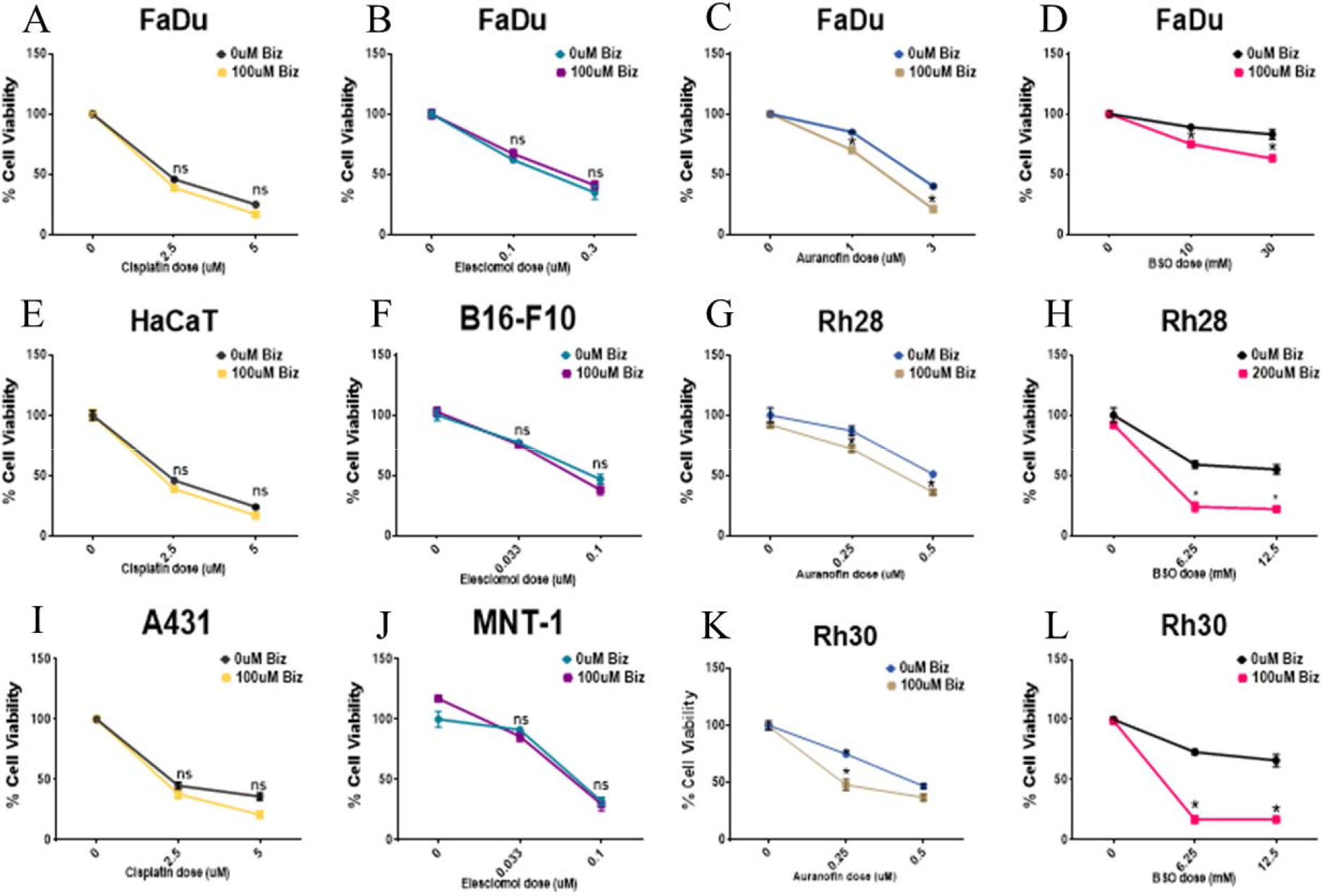
Combination treatment with bizine and anticancer drugs: (A, E, I) Cell viability in FaDu, HaCaT and A-431 cells treated with bizine and cisplatin for 24 hours. (B, F and J) Cell viability in FaDu, B16-F10 and MNT1 cells treated with bizine and cisplatin for 24 hours. (C,G, K) Cell viability in FaDu, Rh28 and Rh30 cells were treated with both bizine and auranofin for 24 hours. (D, H, L) Cell viability in FaDu, Rh28 and Rh30 cells were treated with bizine and BSO for 24 hours. Cell viability was assessed using cell titer blue assay and is expressed as a percentage of control. Points represent mean ± SD (n=3) *: p<0.05.

## Discussion and Conclusion

KDM1A inhibitors have been used in clinical trials as a single anticancer agent with limited success(30,31). Repurposing KDM1A inhibitors as ROS sensitizers identifies a new role for KDM1A as an anticancer redox target. The current studies focus on this novel approach of targeting KDM1A to sensitize cells to exogenous agents that modulate ROS levels to improve anticancer efficacy. Pharmacologic KDM1A inhibition with bizine and phenelzine (Figure 1A-D) and downregulation of KDM1A expression sensitized HaCaT and FaDu cells (Figure 1E-H) to oxidative stress. This phenotype was accompanied by a rapid increase in cellular ROS within 30 min of hydrogen peroxide exposure (Figure 2A-D) and upregulation of ROS responsive genes, KLF-9 and GLRX-1 within 4 hours (Figure 2E-H). Many previous studies involving KDM1A inhibition focused on long term effects with inhibitor treatment ranging from 2-4 days(46–48).

We describe the acute effects of KDM1A inhibition, with effects seen after 1 day of KDM1A inhibition. Previously, in trophoblastic stem cells pharmacological inhibition of KDM1A resulted in an increased ROS generation associated with a senescent phenotype(49). Although we detected a similar increase in ROS, in HaCaT and FaDu cells, only after exogenous oxidative stress and we saw no significant changes in baseline ROS levels, highlighting that the cellular effects of KDM1A inhibition is highly cell context dependent.

Mechanistically, the causes of amplified cellular ROS levels can either be due to increased ROS generation or decreased ROS removal(50). Mitochondrial activity is a major cellular source of ROS(51). KDM1A has been shown to regulate mitochondrial activity in multiple cell types (14,15,38,49,52,53); however, there have been conflicting reports on the effects of KDM1A inhibition on mitochondrial respiration. KDM1A inhibition in white adipocytes, hepatocellular carcinoma cells resulted in increased phosphorylation (15,52), while in brown adipocytes and mouse embryonic fibroblasts the opposite was observed(14,54). In our model, we showed that KDM1A inhibition resulted in decreased basal respiration coupled with reduced ATP generation (Figure 3A-C). These findings indicate that the mitochondrial effect of KDM1A inhibition is both context and cell type dependent. In the cancer cells used in these studies, KDM1A inhibition increased ROS sensitivity without enhancing mitochondrial activity. This is in line with suggestions that the opposing effects of KDM1A inhibition on mitochondrial respiration can be attributed to the differentiation status of cells, as cancer cells frequently exhibit attributes of undifferentiated cells and may even display phenotypic traits of stem cells[54,55]. Given that KDM1A inhibition did not result in an increase in mitochondrial activity and baseline levels of reactive oxygen species (ROS) were not elevated in the absence of an exogenous ROS source, it can be inferred that the inhibition of KDM1A impedes the cell’s capacity to sustain its redox balance when exposed to external ROS stress.

NAC, a synthetic precursor of glutathione, was capable of salvaging cells from bizine- enhanced toxicity (Figure 2Q-R), and GLRX-1 was upregulated by bizine under oxidative stress (Figure 2I and O). KDM1A inhibition with bizine or KDM1A siRNA reduced glutathione levels (Figure 3C). Our data indicate that KDM1A, at least in part, regulates ROS response by maintaining normal physiological levels of glutathione. KDM1A’s effects on metabolic reprogramming in cancer are mediated via HIF-1A (16,39,53,55,56). HIF-1A plays a major role in regulating mitochondrial activity and maintaining levels of antioxidants, including glutathione(57). Our data shows that KDM1A inhibition results in impaired induction of HIF-1A after oxidative stress. Stable downregulation of HIF-1A using shRNA resulted in increased sensitivity to hydrogen peroxide, which was not further increased after bizine treatment (Figure 4F). These results suggest that the effects of bizine on cell viability after oxidative stress are mediated by HIF-1A. The overexpression of constitutively active HIF-1A mitigated the bizine- induced reduction in cell viability but could not completely block bizine from reducing cell viability after hydrogen peroxide treatment. The relationship between KDM1A and HIF-1A has almost exclusively been described under hypoxic conditions (16,39,53,55,56). Our study shows for the first time that KDM1A regulates HIF-1A under oxidative stress, and KDM1A activity modifies HIF-1A mediated antioxidant responses independent of hypoxia.

Our data has so far described the effects of KDM1A on non-histone protein, HIF1A and the impact on cellular metabolism. The impact of KDM1A on non-histone proteins in ROS regulation is one possible mechanism of action; however, we cannot rule out that part of the effects of KDM1A could be due to epigenetic modifications. One such epigenetic target identified in literature was sirtuin 4 (SIRT4) in stem cells. SIRT4 is a mitochondrial protein deacetylase that is important for glutamine anaplerosis and, thus, indirectly regulates levels of glutathione in the cells(58). SIRT4 was shown to mediate the increase in ROS in stem cells after KDM1A inhibition(49). It will be interesting to determine the role of SIRT4 in our and other non-stem cell systems. Moreover, other sirtuins have been linked to HIF-1A regulation. For example, SIRT1 has been shown through deacetylation to increase the stability of HIF-1A under hypoxia(59,60), while SIRT3 suppresses HIF-1A and attenuates transcriptional activity(61).

Additionally, it has been shown that overexpression of KDM1A in cardiac fibroblasts reduced levels of ROS in the cell (62) Recently KDM1A has also been observed to associate with KEAP1 to regulate NRF2 antioxidant gene expression (63) Therefore, KDM1A could play both transcriptional and post-translational roles in the regulation of cellular ROS; future experiments to assess changes in promoter methylation in genes responsible for the metabolism and antioxidant response after KDM1A inhibition will help elucidate this.

KDM1A is significantly upregulated in some cancers, and KDM1A overexpression has been correlated with unfavorable prognostic outcomes (25–29). KDM1A inhibitors have been tested in clinical trials but have had limited success. Combining KDM1A inhibitors with other chemotherapies has shown some promise in acute myeloid leukemia, rhabdomyosarcoma, and breast cancer in vitro(30,31). We have also shown previously that treatment with KDM1A inhibitor greatly improved the efficacy of aminolaevulinic acid (5-ALA) or HPPH (2-[1- hexyloxyethyl]-2-devinyl pyropheophorbide-a) sodium photodynamic therapy in skin cancer cells[32].In our study, we show that combining KDM1A inhibitors with ROS-inducing cancer therapies may pose an effective anticancer strategy. We tested multiple drugs which have been reported in the literature to increase ROS, including cisplatin, elesclomol, auranofin, and BSO (Figure 5) in relevant *in vitro* models, including squamous cell carcinoma, melanoma, and rhabdomyosarcoma in combination with bizine. Bizine only enhanced cell death in combination with BSO and auranofin and was incapable of increasing cisplatin or elesclomol-induced death. While hydrogen peroxide is understood to be a source of ROS with cellular effects mediated by its redox effects, most ROS-inducing chemotherapeutic agents generate ROS as a by-product of other toxic effects(64). The cytotoxicity of the compounds that were unsuccessful may not be primarily mediated by ROS upregulation. Alternatively, a secondary change in cells committed to death, or the ROS produced by these compounds is confined to cell compartments where the inhibition of KDM1A does not disrupt the cell’s antioxidant defenses, are also possible. This latter possibility is supported by the fact that elesclomol and arsenic trioxide specifically target the mitochondria and primarily generate mitochondrial superoxides(42,65). These findings indicate that the source and localization of ROS should be considered when selecting candidates for combination treatment with bizine.

Auranofin and BSO primarily target the antioxidant machinery and are used in clinical trials for the treatment of rhabdomyosarcoma(43). Auranofin abrogates the thioredoxin system(66) and BSO inhibits glutathione production(45). BSO and auranofin were effective in enhancing cell death in combination with bizine in FaDu and rhabdomyosarcoma cells (Figure 6). As KDM1A inhibition decreased cellular levels of glutathione, it is not surprising that combining bizine with inhibitors of endogenous antioxidants enhanced cell death. Our data show for the first time that dual targeting of KDM1A and antioxidant systems is a possible new treatment combination for cancers susceptible to ROS imbalance. Our findings have added to our understanding of the complex role KDM1A plays in cancer and opened new potential avenues for developing KDM1A inhibitor-based cancer therapies.

**Figure 6:**
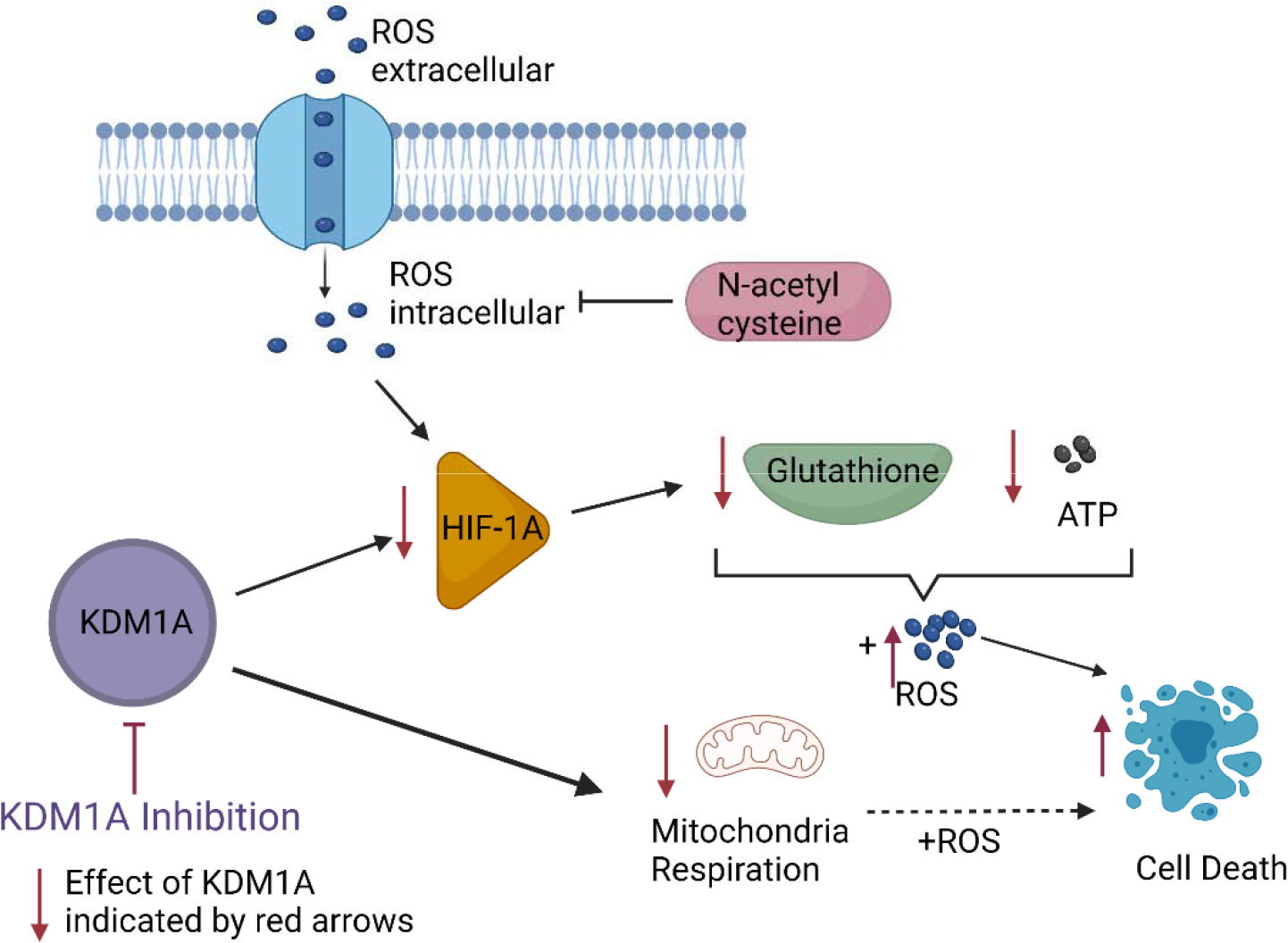
**Mechanism of action of KDM1A inhibition in ROS regulation**

## Acknowledgements

This research was funded by the Roswell Park Alliance Foundation (PI: Paragh) and NCI grant P30 CA016056 involving the use of Roswell Park Comprehensive Cancer Center’s Shared Resources: Flow and Image Cytometry and Immune Analysis.

## Bibliography

1. Szatrowski TP, Nathan CF. Production of Large Amounts of Hydrogen Peroxide by Human Tumor Cells. Cancer Res. 1991;

2. Wu WS. The signaling mechanism of ROS in tumor progression. Cancer and Metastasis Reviews. 2006.

3. Timblin CR, Janssen YWM, Mossman BT. Transcriptional Activation of the Proto-oncogene c-jun by Asbestos and H2O2 Is Directly Related to Increased Proliferation and Transformation of Tracheal Epithelial Cells. Cancer Res. 1995;

4. Lee SF, Huang YT, Wu WS, Lin JK. Induction of c-jun protooncogene expression by hydrogen peroxide through hydroxyl radical generation and p60(src) tyrosine kinase activation. Free Radic Biol Med. 1996;

5. Li DWC, Spector A. Hydrogen peroxide-induced expression of the proto-oncogenes, c-jun, c-fos and c-myc in rabbit lens epithelial cells. Mol Cell Biochem. 1997;

6. Sabri A, Byron KL, Samarel AM, Bell J, Lucchesi PA. Hydrogen peroxide activates mitogen-activated protein kinases and Na+- H+ exchange in neonatal rat cardiac myocytes. Circ Res. 1998;

7. Son Y, Kim S, Chung HT, Pae HO. Reactive oxygen species in the activation of MAP kinases. In: Methods in Enzymology. 2013.

8. Meves A, Stock SN, Beyerle A, Pittelkow MR, Peus D. H2O2 mediates oxidative stress-induced epidermal growth factor receptor phosphorylation. Toxicol Lett. 2001;122(3):205– 14.

9. Zwang Y, Yarden Y. p38 MAP kinase mediates stress-induced internalization of EGFR: Implications for cancer chemotherapy. EMBO J. 2006;

10. Liu Y, Li Q, Zhou L, Xie N, Nice EC, Zhang H, et al. Cancer drug resistance: redox resetting renders a way. Oncotarget. 2016;7(27):42740–61.

11. Harris IS, Treloar AE, Inoue S, Sasaki M, Gorrini C, Lee KC, et al. Glutathione and Thioredoxin Antioxidant Pathways Synergize to Drive Cancer Initiation and Progression. Cancer Cell. 2015;

12. Stavropoulos P, Blobel G, Hoelz A. Crystal structure and mechanism of human lysine-specific demethylase-1. Nat Struct Mol Biol. 2006;13(7):626–32.

13. Nicholson TB, Chen T. LSD1 demethylates histone and non-histone proteins. Epigenetics. 2009;4(3):129–32.

14. Duteil D, Tosic M, Lausecker F, Nenseth HZ, Müller JM, Urban S, et al. Lsd1 Ablation Triggers Metabolic Reprogramming of Brown Adipose Tissue. Cell Rep. 2016;17(4):1008–21.

15. Duteil D, Metzger E, Willmann D, Karagianni P, Friedrichs N, Greschik H, et al. LSD1 promotes oxidative metabolism of white adipose tissue. Nat Commun. 2014;5:4093.

16. Baek SH, Kim K Il. Regulation of HIF-1α stability by lysine methylation. BMB Rep. 2016;49(5):245–6.

17. Huang J, Sengupta R, Espejo AB, Lee MG, Dorsey JA, Richter M, et al. p53 is regulated by the lysine demethylase LSD1. Nature. 2007;449(7158):105–8.

18. Tsai W-W, Nguyen TT, Shi Y, Barton MC. p53-targeted LSD1 functions in repression of chromatin structure and transcription in vivo. Mol Cell Biol. 2008;28(17):5139–46.

19. Peng B, Wang J, Hu Y, Zhao H, Hou W, Zhao H, et al. Modulation of LSD1 phosphorylation by CK2/WIP1 regulates RNF168-dependent 53BP1 recruitment in response to DNA damage. Nucleic Acids Res. 2015;43(12):5936–47.

20. Kontaki H, Talianidis I. Lysine methylation regulates E2F1-induced cell death. Mol Cell. 2010;39(1):152–60.

21. Xie Q, Bai Y, Wu J, Sun Y, Wang Y, Zhang Y, et al. Methylation-mediated regulation of E2F1 in DNA damage-induced cell death. J Recept Signal Transduct Res. 2011;31(2):139–46.

22. He Y, Zhao Y, Wang L, Bohrer LR, Pan Y, Wang L, et al. LSD1 promotes S-phase entry and tumorigenesis via chromatin co-occupation with E2F1 and selective H3K9 demethylation. Oncogene. 2018;37(4):534–43.

23. Amente S, Milazzo G, Sorrentino MC, Ambrosio S, Di Palo G, Lania L, et al. Lysine-specific demethylase (LSD1/KDM1A) and MYCN cooperatively repress tumor suppressor genes in neuroblastoma. Oncotarget. 2015;6(16):14572–83.

24. Ambrosio S, Amente S, Saccà CD, Capasso M, Calogero RA, Lania L, et al. LSD1 mediates MYCN control of epithelial-mesenchymal transition through silencing of metastatic suppressor NDRG1 gene. Oncotarget. 2017;8(3):3854–69.

25. Zhao Z-K, Yu H-F, Wang D-R, Dong P, Chen L, Wu W-G, et al. Overexpression of lysine specific demethylase 1 predicts worse prognosis in primary hepatocellular carcinoma patients. World J Gastroenterol. 2012;18(45):6651–6.

26. Lv T, Yuan D, Miao X, Lv Y, Zhan P, Shen X, et al. Over-Expression of LSD1 Promotes Proliferation, Migration and Invasion in Non-Small Cell Lung Cancer. PLoS One. 2012;7(4):e35065.

27. Jie D, Zhongmin Z, Guoqing L, Sheng L, Yi Z, Jing W, et al. Positive expression of LSD1 and negative expression of E-cadherin correlate with metastasis and poor prognosis of colon cancer. Dig Dis Sci. 2013;58(6):1581–9.

28. Nagasawa S, Sedukhina AS, Nakagawa Y, Maeda I, Kubota M, Ohnuma S, et al. LSD1 overexpression is associated with poor prognosis in basal-like breast cancer, and sensitivity to PARP inhibition. PLoS One. 2015;10(2):e0118002.

29. Lian SX, Shao YB, Liu HB, He JY, Lu WQ, Zhang Y, et al. Lysine-specific demethylase 1 promotes tumorigenesis and predicts prognosis in gallbladder cancer. Oncotarget. 2015;6(32):33065–76.

30. Haydn T, Metzger E, Schuele R, Fulda S. Concomitant epigenetic targeting of LSD1 and HDAC synergistically induces mitochondrial apoptosis in rhabdomyosarcoma cells. Cell Death Dis. 2017;

31. Duan YC, Ma YC, Qin WP, Ding LN, Zheng YC, Zhu YL, et al. Design and synthesis of tranylcypromine derivatives as novel LSD1/HDACs dual inhibitors for cancer treatment. Eur J Med Chem. 2017;

32. Mudambi S, Fitzgerald M, Pera P, Washington D, Chamberlain S, Fidrus E, et al. KDM1A inhibition increases UVA toxicity and enhances photodynamic therapy efficacy. Photodermatol Photoimmunol Photomed. 2023 May;39(3):226–34.

33. Bagati A, Bianchi-Smiraglia A, Moparthy S, Kolesnikova K, Fink EE, Kolesnikova M, et al. FOXQ1 controls the induced differentiation of melanocytic cells. Cell Death Differ. 2018;25(6):1040–9.

34. Bao L, Chen Y, Lai H-T, Wu S-Y, Wang JE, Hatanpaa KJ, et al. Methylation of hypoxia-inducible factor (HIF)-1α by G9a/GLP inhibits HIF-1 transcriptional activity and cell migration. Nucleic Acids Res. 2018 Jul;46(13):6576–91.

35. Anathy V, Aesif SW, Guala AS, Havermans M, Reynaert NL, Ho YS, et al. Redox amplification of apoptosis by caspase-dependent cleavage of glutaredoxin 1 and S- glutathionylation of Fas. J Cell Biol. 2009;

36. Sauer H, Wartenberg M, Hescheler J. Reactive oxygen species as intracellular messengers during cell growth and differentiation. Cell Physiol Biochem. 2001;11(4):173– 86.

37. Samanta D, Semenza GL. Maintenance of redox homeostasis by hypoxia-inducible factors. Redox Biology. 2017.

38. Kosumi K, Baba Y, Sakamoto A, Ishimoto T, Harada K, Nakamura K, et al. Lysine-specific demethylase-1 contributes to malignant behavior by regulation of invasive activity and metabolic shift in esophageal cancer. Int J Cancer. 2016;138(2):428–39.

39. Qin Y, Zhu W, Xu W, Zhang B, Shi S, Ji S, et al. LSD1 sustains pancreatic cancer growth via maintaining HIF1α-dependent glycolytic process. Cancer Lett. 2014;347(2):225–32.

40. Ketscher A, Jilg CA, Willmann D, Hummel B, Imhof A, Rüsseler V, et al. LSD1 controls metastasis of androgen-independent prostate cancer cells through PXN and LPAR6. Oncogenesis. 2014;3:e120.

41. Choi Y-M, Kim H-K, Shim W, Anwar MA, Kwon J-W, Kwon H-K, et al. Mechanism of Cisplatin-Induced Cytotoxicity Is Correlated to Impaired Metabolism Due to Mitochondrial ROS Generation. PLoS One. 2015;10(8):e0135083.

42. Kirshner JR, He S, Balasubramanyam V, Kepros J, Yang C-Y, Zhang M, et al. Elesclomol induces cancer cell apoptosis through oxidative stress. Mol Cancer Ther. 2008;7(8):2319–27.

43. Habermann KJ, Grünewald L, van Wijk S, Fulda S. Targeting redox homeostasis in rhabdomyosarcoma cells: GSH-depleting agents enhance auranofin-induced cell death. Cell Death Dis. 2017;

44. Roder C, Thomson MJ. Auranofin: Repurposing an Old Drug for a Golden New Age. Drugs R D. 2015;15(1):13–20.

45. Tagde A, Singh H, Kang MH, Reynolds CP. The glutathione synthesis inhibitor buthionine sulfoximine synergistically enhanced melphalan activity against preclinical models of multiple myeloma. Blood Cancer J. 2014;4:e229.

46. Maes T, Mascaró C, Tirapu I, Estiarte A, Ciceri F, Lunardi S, et al. ORY-1001, a Potent and Selective Covalent KDM1A Inhibitor, for the Treatment of Acute Leukemia. Cancer Cell. 2018;

47. Sehrawat A, Gao L, Wang Y, Bankhead A, McWeeney SK, King CJ, et al. LSD1 activates a lethal prostate cancer gene network independently of its demethylase function. Proc Natl Acad Sci U S A. 2018;

48. Verigos J, Karakaidos P, Kordias D, Papoudou-Bai A, Evangelou Z, Harissis H V., et al. The histone demethylase LSD1/KDM1A mediates chemoresistance in breast cancer via regulation of a stem cell program. Cancers (Basel). 2019;

49. Castex J, Willmann D, Kanouni T, Arrigoni L, Li Y, Friedrich M, et al. Inactivation of Lsd1 triggers senescence in trophoblast stem cells by induction of Sirt4. Cell Death Dis. 2017;8(2):e2631.

50. Panieri E, Santoro MM. ROS homeostasis and metabolism: a dangerous liason in cancer cells. Cell Death Dis. 2016;7(6):e2253.

51. Ciccarese F, Ciminale V. Escaping Death: Mitochondrial Redox Homeostasis in Cancer Cells. Front Oncol. 2017;7:117.

52. Sakamoto A, Hino S, Nagaoka K, Anan K, Takase R, Matsumori H, et al. Lysine demethylase LSD1 coordinates glycolytic and mitochondrial metabolism in hepatocellular carcinoma cells. Cancer Res. 2015;75(7):1445–56.

53. Sakamoto A, Hino S, Nagaoka K, Anan K, Takase R, Matsumori H, et al. Lysine Demethylase LSD1 Coordinates Glycolytic and Mitochondrial Metabolism in Hepatocellular Carcinoma Cells. Cancer Res. 2015;75(7):1445–56.

54. Sun H, Liang L, Li Y, Feng C, Li L, Zhang Y, et al. Lysine-specific histone demethylase 1 inhibition promotes reprogramming by facilitating the expression of exogenous transcriptional factors and metabolic switch. Sci Rep. 2016;6:30903.

55. Lee J-Y, Park J-H, Choi H-J, Won H-Y, Joo H-S, Shin D-H, et al. LSD1 demethylates HIF1α to inhibit hydroxylation and ubiquitin-mediated degradation in tumor angiogenesis. Oncogene. 2017;36(39):5512–21.

56. Yang S-J, Park YS, Cho JH, Moon B, An H-J, Lee JY, et al. Regulation of hypoxia responses by flavin adenine dinucleotide-dependent modulation of HIF-1α protein stability. EMBO J. 2017;36(8):1011–28.

57. Krüger A, Grüning N-M, Wamelink MMC, Kerick M, Kirpy A, Parkhomchuk D, et al. The pentose phosphate pathway is a metabolic redox sensor and regulates transcription during the antioxidant response. Antioxid Redox Signal. 2011;15(2):311–24.

58. Fernandez-Marcos PJ, Serrano M. Sirt4: the glutamine gatekeeper. Cancer Cell. 2013;23(4):427–8.

59. Lim J-H, Lee Y-M, Chun Y-S, Chen J, Kim J-E, Park J-W. Sirtuin 1 modulates cellular responses to hypoxia by deacetylating hypoxia-inducible factor 1alpha. Mol Cell. 2010;38(6):864–78.

60. Laemmle A, Lechleiter A, Roh V, Schwarz C, Portmann S, Furer C, et al. Inhibition of SIRT1 impairs the accumulation and transcriptional activity of HIF-1α protein under hypoxic conditions. PLoS One. 2012;7(3):e33433.

61. Finley LWS, Carracedo A, Lee J, Souza A, Egia A, Zhang J, et al. SIRT3 opposes reprogramming of cancer cell metabolism through HIF1α destabilization. Cancer Cell. 2011;19(3):416–28.

62. Cao Y, Dong Z, Yang D, Ma X, Wang X. LSD1 regulates the expressions of core cardiogenic transcription factors and cardiac genes in oxygen and glucose deprivation injured mice fibroblasts in vitro. Exp Cell Res. 2022;418(1):113228.

63. Lin C-Y, Chang C-B, Wu R-C, Chao A, Lee Y-S, Tsai C-N, et al. Glucose Activates Lysine-Specific Demethylase 1 through the KEAP1/p62 Pathway. Antioxidants. 2021;10(12).

64. Yang H, Villani RM, Wang H, Simpson MJ, Roberts MS, Tang M, et al. The role of cellular reactive oxygen species in cancer chemotherapy. J Exp Clin Cancer Res. 2018 Nov;37(1):266.

65. Woo SH, Park IC, Park MJ, Lee HC, Lee SJ, Chun YJ, et al. Arsenic trioxide induces apoptosis through a reactive oxygen species-dependent pathway and loss of mitochondrial membrane potential in HeLa cells. Int J Oncol. 2002;

66. Marzano C, Gandin V, Folda A, Scutari G, Bindoli A, Rigobello MP. Inhibition of thioredoxin reductase by auranofin induces apoptosis in cisplatin-resistant human ovarian cancer cells. Free Radic Biol Med. 2007;42(6):872–81.

