## Supplemental Figure 1 for "Dual targeting of KDM1A and antioxidants is an effective anticancer strategy"

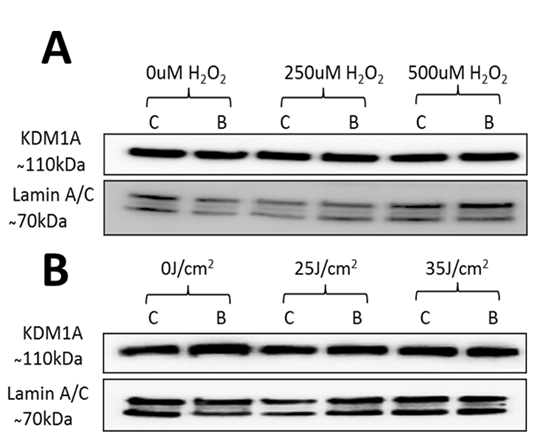

Figure S1: KDM1A protein regulation after hydrogen peroxide exposure in FaDu cells -Western blot showing KDM1A protein expression in FaDu cells after 24 hours of bizine treatment and 4 hours of hydrogen peroxide.
