## Supplemental Figure 2 for "Dual targeting of KDM1A and antioxidants is an effective anticancer strategy"

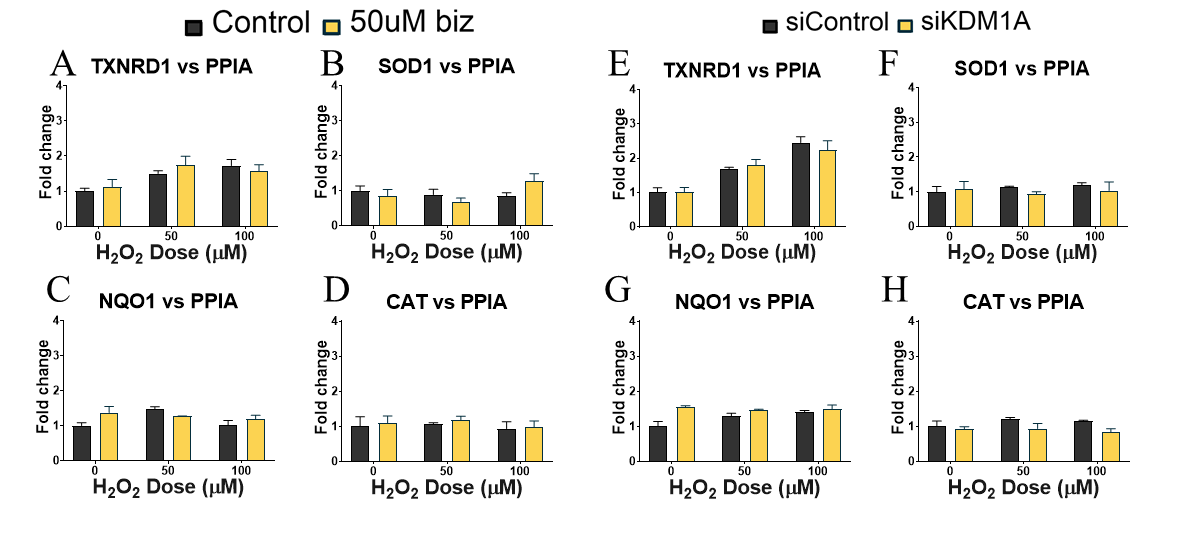

Figure S2: Gene expression of ROS regulated genes (TXNRD1 SOD1, NQO1 and CAT) after 24 hours of 50µM bizine treatment in FaDu cells (A-D) or in KDM1A downregulated FaDu cells (E-H).
