## Supplemental Figure 3 for "Dual targeting of KDM1A and antioxidants is an effective anticancer strategy"

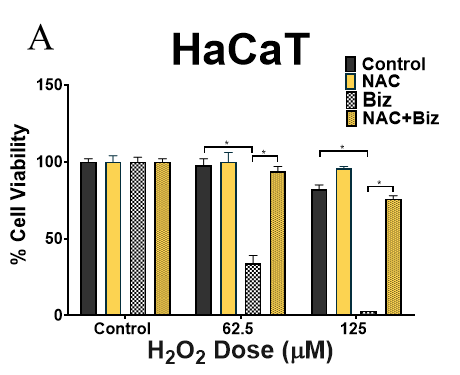

Figure S3: Cell viability in HaCaT cells after 18 hours of 150uM of bizine and 10mM NAC treatment and hydrogen peroxide exposure Cell viability was assessed using cell titer blue assay and is expressed as a percentage of control. Bars represent mean ± SD (n=3). *: p<0.05 calculated using student t-test.
