## Supplemental Figure 4 for "Dual targeting of KDM1A and antioxidants is an effective anticancer strategy"

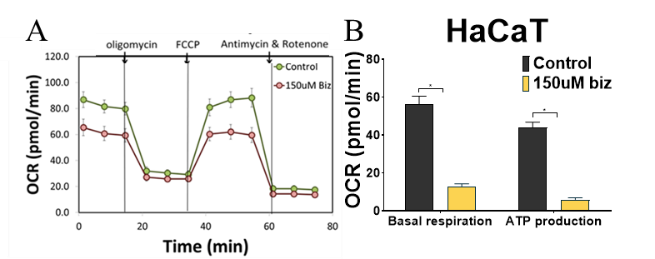

Figure S4: (A) MitoStress test using seahorse assay in HaCaT cells after bizine treatment. (B) Graphs quantifying basal respiration and ATP generation in bizine treated HaCaT cells.
